# Cytoskeletal vimentin regulates cell size and autophagy through mTORC1 signaling

**DOI:** 10.1101/2021.04.19.440145

**Authors:** Ponnuswamy Mohanasundaram, Leila S Coelho Rato, Mayank Modi, Marta Urbanska, Franziska Lautenschläger, Fang Cheng, John E Eriksson

## Abstract

The nutrient-activated mTORC1 (mechanistic target of rapamycin kinase complex 1) signaling pathway determines cell size by controlling mRNA translation, ribosome biogenesis, protein synthesis, and autophagy. Here we show that vimentin, a cytoskeletal intermediate filament protein that we know to be important for wound healing and cancer progression, determines cell size through mTORC1 signaling, an effect that is also manifested at the organism level in mice. We found that vimentin maintains normal cell size by supporting mTORC1 activation and through inhibition of autophagic flux. This regulation is manifested at all levels of downstream target activation and regulation of protein synthesis. We show that vimentin controls mTORC1 mobility by allowing access to lysosomes. Vimentin inhibits the autophagic flux in normal fibroblasts even under starved conditions, indicating a growth factor-independent inhibition of autophagy at the level of mTORC1. Our findings demonstrate that vimentin couples cell size signaling and autophagy with the biomechanic, sensing, and kinetic functions of the cytoskeleton.

---

Cell size regulation is intricately related to nutrient availability and the rate by which macromolecules crucial for cell functions and buildup are synthesized (1). mTORC1 acts as central signaling hub in regulating cell size and metabolism by sensing and coordinating stimuli derived from nutrients, energy, stress, and growth factors(2). In the presence of amino acids, Rag GTPases recruit mTORC1 to the lysosomal membrane where it is activated by Rheb(3–5). This allows mTORC1 to promote protein synthesis by controlling two downstream effectors, 4-EBP1 and p70S6K(6) and by inhibiting autophagy via phosphorylation ULK1, a known promoter of autophagy(7–9).

Vimentin, an intermediate filament (IFs) protein, has been reported to act as signaling scaffold in key cellular processes required to maintain tissue integrity and facilitate tissue repair(10,11). These include cell migration, adhesion, proliferation and invasion(12). In this respect, vimentin is a well-established marker for epithelial to mesenchymal transition (EMT), a crucial step in wound healing and metastasis(13). Vimentin-deficient (Vim -/-) mice exhibit delayed wound healing due to defects in EMT signaling, cell migration, and cell proliferation(14). Furthermore, vimentin regulates autophagy by forming a complex with the autophagy regulator Beclin and the adaptor protein 14-3-3(15). Another link between cell size and vimentin was provided by a recent study showing that Vim -/- mice have deficient accumulation of body fat(16) and by the first report on a human vimentin mutation leading to lipodystrophy(17, 41). Keratin 17, another member of the IF family, was implicated in cell size signaling, as mouse-derived skin keratinocytes lacking keratin 17 are smaller than corresponding WT cells and display lower AKT/mTOR activation(18). Related to all these studies, we have observed that cell size regulation is coupled to vimentin, both in terms of enlargement and reduction.

When reporting that vimentin has a role in fibroblast proliferation and in EMT(14,19), we observed that not only do Vim -/- fibroblasts grow slower, but they were also significantly smaller (Extended Data Video 1; https://youtu.be/O_mbLL0i10). Correspondingly, when examining the Vim-/- mice in greater detail, we observed that they were leaner, as reflected in remarkable reductions in weight, body-mass index, as well as Lee’s index (Fig. 1a–c), preliminary indicated in a recent report (16). These mice also had markedly lower fat content (Fig. 1d), and smaller size of adipocytes (Fig. 1e). Likewise, some of the organs (heart and kidney) were also smaller in size (Fig. 1f). As all these effects point towards a potential disturbance in cell size regulation, we quantified in detail the effects of vimentin on cell size. We found that Vim -/- MEFs are markedly smaller than WT MEFs (Fig. 2a, b). This effect was seen in cells in their active growth phase and cells in confluent state (Extended Data Video 1; https://youtu.be/O_mbLL0i10), indicating that this effect is not coupled to cell spreading. To eliminate the effect of cell spreading, we next measured the cell volume of trypsinized MEFs and found that the volume of MEFs Vim -/- is significantly lower than MEFs WT (Fig. 2c, d). To minimize cell cycle-dependent size variation we employed thymidine-induced cell cycle arrest, which revealed an even more pronounced cell volume reduction in the Vim -/- MEFs as compared to the WT MEFs, demonstrating that the size effect on cell volume is cell cycle-independent (Fig. 2e). Moreover, we could rescue the cell volume by reintroducing WT vimentin into Vim -/- MEFs (Fig. 2f). To ensure that these effects were not specific for the immortalized MEFs, we assessed the size of primary MEFs isolated from WT and Vim -/- mice, which showed a similar size reduction in the Vim -/- primary MEFs (Fig. 2g). We also investigated the size of primary bone marrow-derived dendritic cells (BMDCs) using a high-throughput microfluidics-based method (20), to compare the results acquired from adherent fibroblasts with a non-adherent and spherical cell type. These cells possess a less extensive cytoskeleton, which implies that possible structural effects from cytoskeletal elements other than vimentin are minimized. Furthermore, we used a microfluidic method to image the cells not only in the initial state, but also upon deformation in a narrow constriction of a microfluidic channel, where cytoskeletal structures are strained. Consistently with the above results, BMDCs lacking vimentin showed decreased size in both normal and deformed state (Extended Data Fig. 1a, b). These results emphasize the general nature of the vimentin-dependent effect on cell size.

**Figure. 1.**
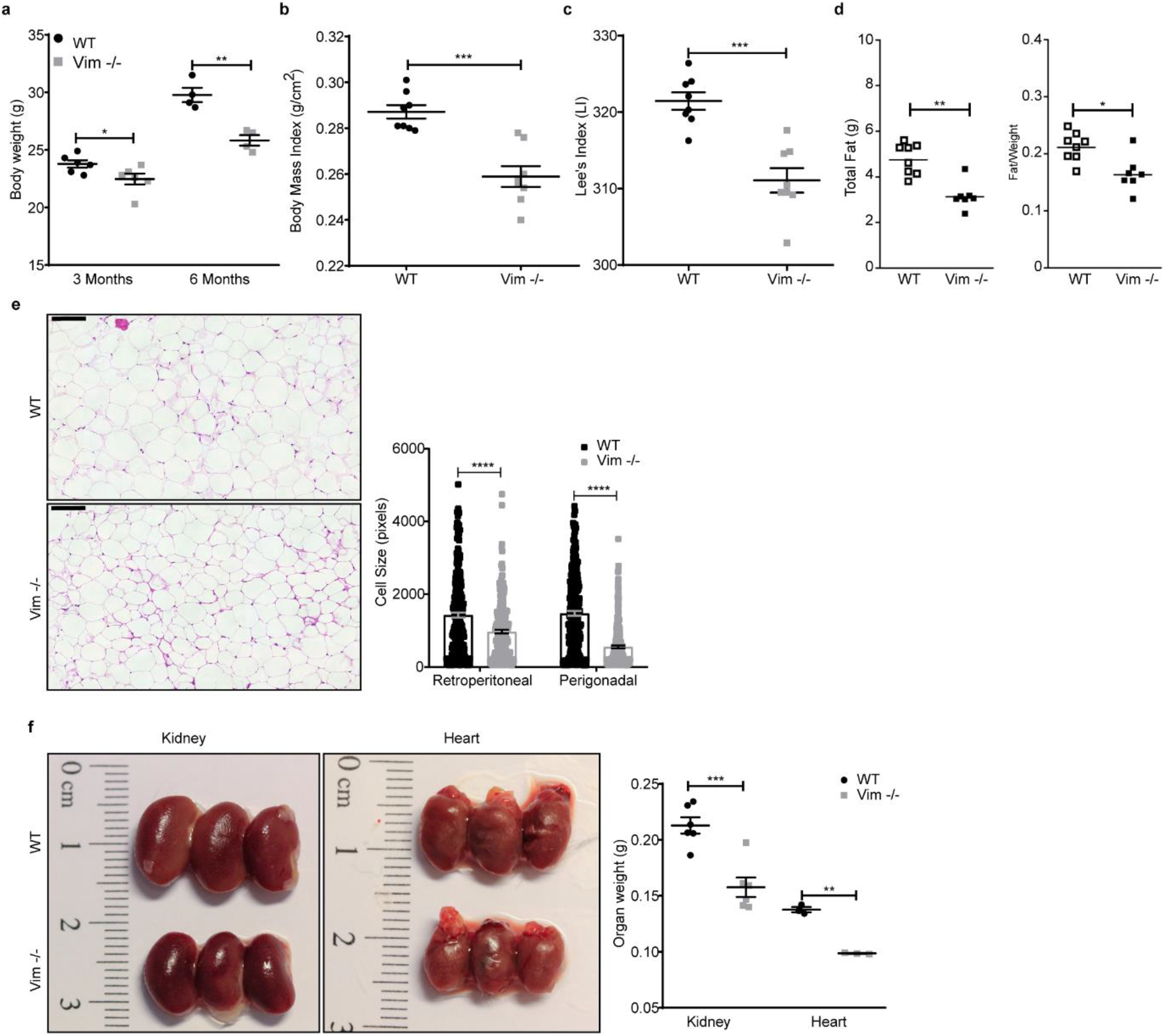
Vim -/- mice are smaller than WT. **a.** Body weight of WT and Vim -/- male mice of 3 (n=6) and 6 months old (n=4). **b.** BMI and **c.** Lee’s index were calculated for 6 months old mice. **d.** Total fat mass determined by EcoMRI and ratio of fat/weight of 6 months old male mice. **e.** Micrograph of adipocytes from 3 months old WT and Vim -/- male mice. The size of retroperitoneal and perigonadal adipocytes was quantified using Fiji. **f.** Kidney (n=6) and heart (n=3) organ weight of 3 months old male mice. The results are presented in the form of mean ± standard deviation of the mean of the biological replicates. *p<0.05; **p<0.01, ***p<0.001, ****p<0.0001.

**Figure. 2.**
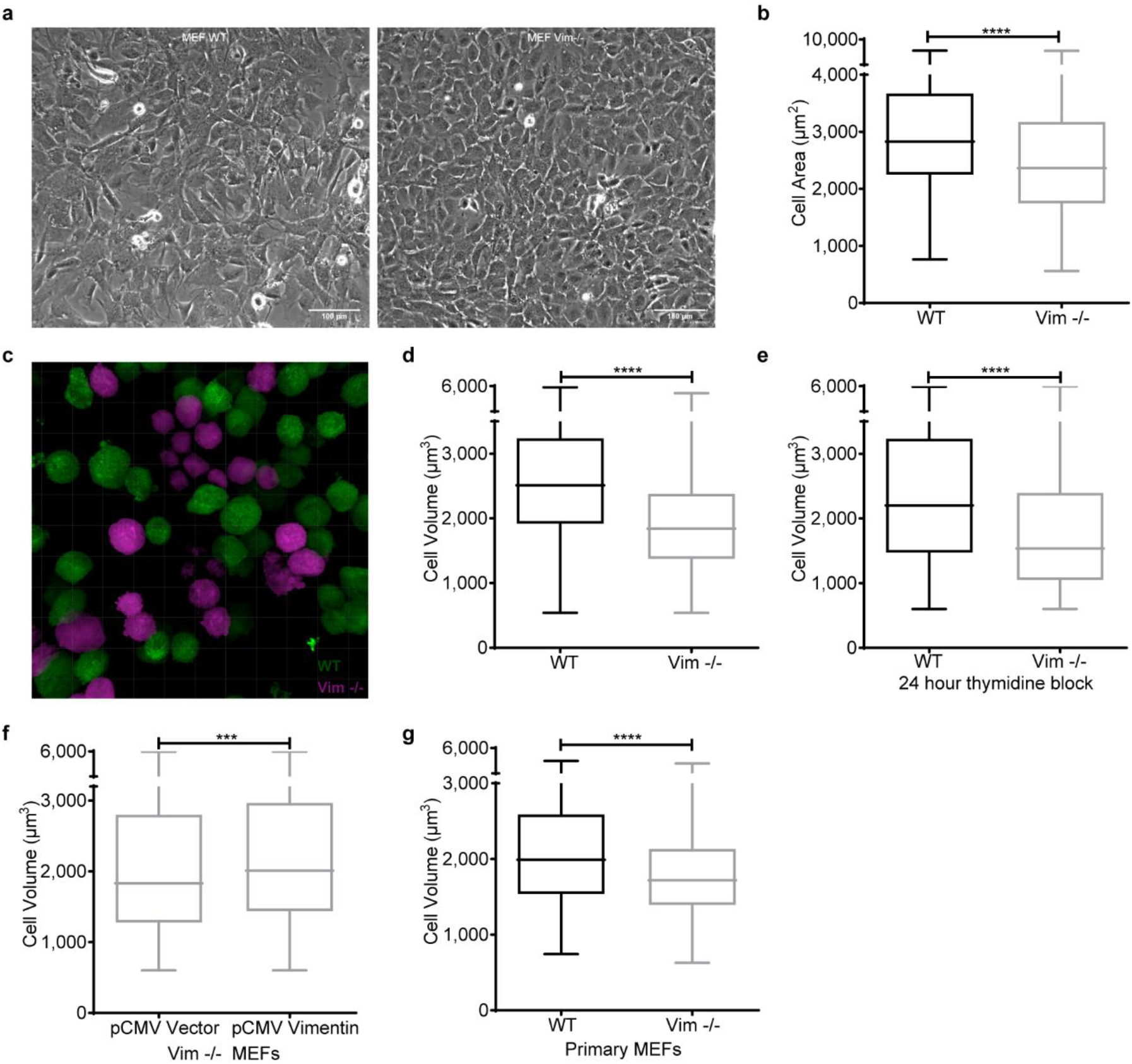
Loss of vimentin reduces cell size. **a.** Phase contrast representative images of WT and Vim -/- MEFs (Scale bar = 100 μm). **b.** Distribution of the cell area obtained from phase contrast images measured using Fiji (n=3). **c.** Micrograph of WT (green) and Vim -/- (magenta) MEFs image made with Imaris software for cell volume analysis. **d.** Cell volume distribution of WT and Vim -/- MEFs obtained from z-stacks from cells in suspension stained with CellTracker (n=3). **e.** Cell volume distribution of WT and Vim -/- cells treated with 1 mM of thymidine for 24 hours (n=3). **f.** Cell volume distribution after transfection of empty vector or pCMV script-vimentin into Vim -/- MEFs. Cells were transfected with 7.5 μg of plasmid and the cell volume analysis was conducted 48 hours post-transfection (n=3). **g.** Cell volume distribution of primary MEFs isolated for WT and Vim -/- mice (n=3). ***p<0.001, ****p<0.0001.

As cell size is tightly regulated by nutrients and growth factors(1), we wanted to assess whether the effect seen on cell size is linked to these stimuli. To this end, cells were serum-starved for two to three days, then transferred to fresh serum-free media, and subsequently stimulated with insulin or fetal calf serum (FCS) for 24 hours. Importantly, we found that serum starvation reduced the size of WT MEFs to the same as Vim -/- MEFs. Stimulation with insulin only, which rules out the influence of other growth factors, increased cell size only in MEFs WT (Fig. 3a). Consistently, when the cells were stimulated with FCS, there was a significant increase in WT MEFs, whereas Vim -/- MEFs were unaffected. Our results show that the reduction in cell size in Vim -/- MEFs is linked to a disruption in insulin-dependent signaling. Thus, the smaller size stems from abrogation of the cellular control mechanisms regulating this attribute and not from structural effects of a missing cytoskeletal component. Since cell size depends on the balance between synthesis and degradation of macromolecules(1), we measured protein amount per cell and found that it was significantly lower in MEFs Vim -/- (Fig. 3b), indicating that the reduction in cell size is directly coupled to the anabolic state of the cells. To link this observation to growth signaling, we measured the level of protein synthesis after insulin stimulation and observed that it was significantly lower in Vim -/- MEFs (Fig. 3c). These results demonstrate that vimentin participates in signaling stimulating cell size. Interestingly, starvation reduced the size of WT MEFs to the size of vimentin-deficient cells, while the size of the Vim-/- MEFs was not affected. This implies that the Vim-/-MEFs behave as if they would be in a compromised nutritional state already without starvation, due to deficient nutrient-coupled signaling.

It is well established that mTORC1 is one of the main pathways regulating cell size and metabolism in response to nutrients, growth factors, and other extracellular cues(21). To investigate the relationship between mTORC1 signaling and the role of vimentin in cell size, we analyzed mTORC1 activity after overnight serum starvation of cells and subsequent stimulation with insulin or FCS for 15 minutes. Intriguingly, we found that already without stimulation, the phosphorylation of mTORC1 downstream targets were lower in Vim -/- MEFs. Strikingly, Vim -/- MEFs stimulated with insulin or FCS showed only negligible mTORC1 activation (Fig. 3d), indicating that the observed faulty cell size signaling is due to a defect in mTORC1 activation. Together, these results show that the cell size reduction in Vim -/- cells is due to abrogated mTORC1 signaling.

**Figure. 3.**
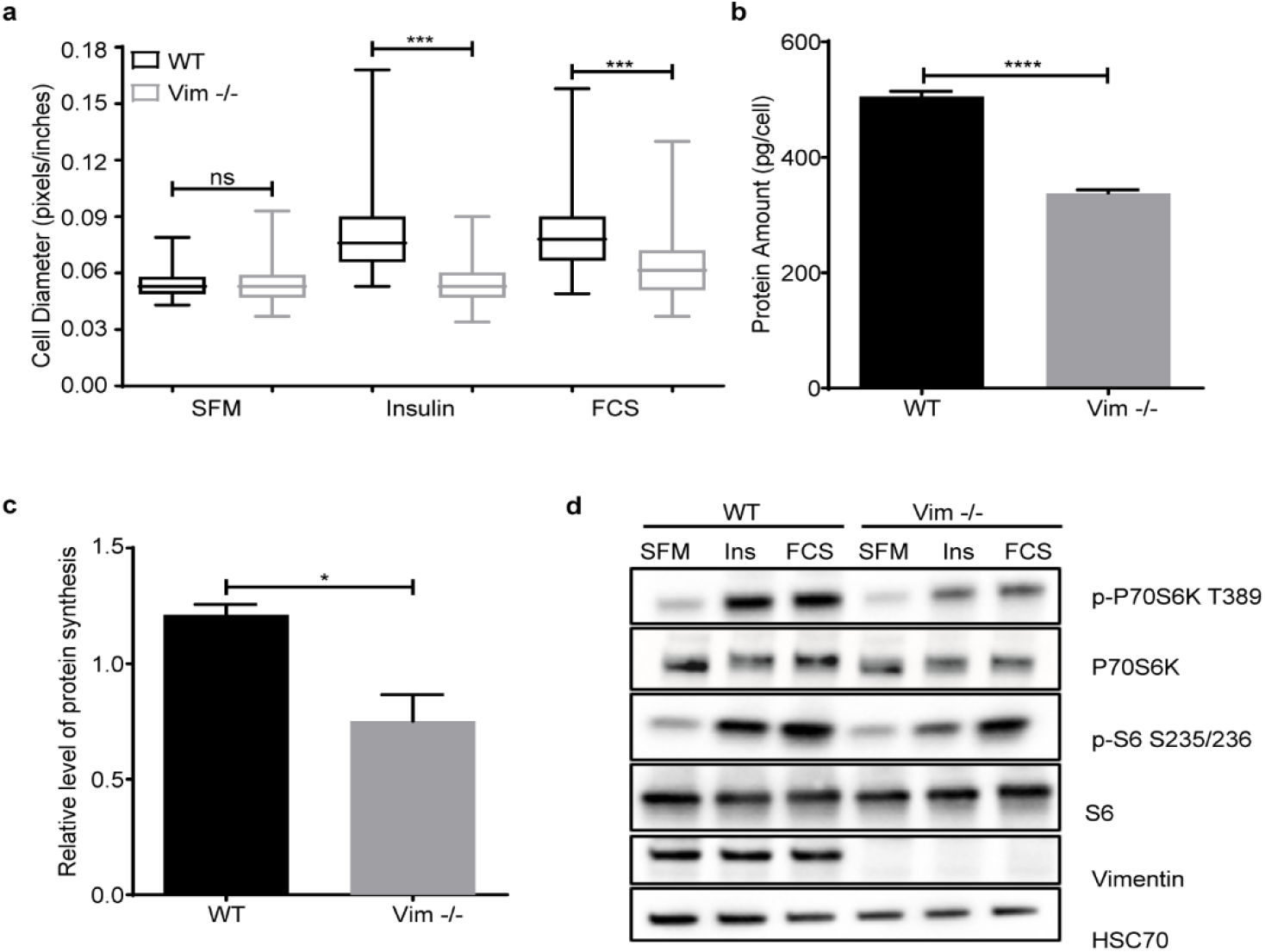
mTORC1 translocation and activation is impaired in Vim -/- MEFs. **a.** Distribution of the cell diameter of WT and Vim -/- MEFs serum-starved for 36 hours and treated with 100 μg/ml insulin or 5 % FCS for 24 hours (n=3). **b.** Protein amount per cell is represented as mean ± SEM of protein concentration. An equal number of cells were lysed, and protein concentration was measured using a commercial BCA kit (n=3). **c**. Protein synthesis is represented as mean ± SEM of WT and Vim -/- MEFs stimulated with insulin for 30 minutes (n=3). Protein synthesis was quantified using the mean fluorescence intensity normalized to the serum-starved treatment. **d.** Western blotting analysis of serum-starved cells (overnight) treated with 100 nM of insulin or 5 % serum for 15 minutes (n=3).*p<0.05, ***p<0.001, ****p<0.0001, ns=non-significant.

Along with insulin and growth factor signaling, mTORC1 is activated by nutrients (6). To understand how the combination of nutrients and insulin signaling modulates vimentin-mediated mTORC1 signaling, we starved the cells for one hour in culture media without amino acids, glucose, and serum, followed by stimulation with amino acids L-glutamine with minimum essential amino acids (EAAs), or EAAs and non-EAAs, glucose and insulin. As Vim -/- MEFs displayed a striking defect in the insulin-mediated stimulation of mTORC1 signaling (Fig. 4a), we first studied what are the differences in the presence of insulin while the nutrient sources are varied. By maximal stimulation of insulin-mediated signaling, we wanted to reveal possible differential effects of variable nutrient sources and, in this way, examine the upstream parts of the insulin signaling pathway. The lack of a difference in AKT phosphorylation (Fig. 4a), implies that vimentin regulates mTORC1 signaling primarily in the downstream parts of the pathway. Importantly, when examining the downstream targets, we saw that when insulin signaling is pushed to the maximum, the downstream signaling is significantly suppressed regardless of the nutrient source (Fig. 4a). This outcome demonstrates that the capacity of insulin signaling to amplify the mTORC1 pathway is always dependent on vimentin, regardless of which additional stimuli is provided by individual nutrients.

**Figure. 4.**
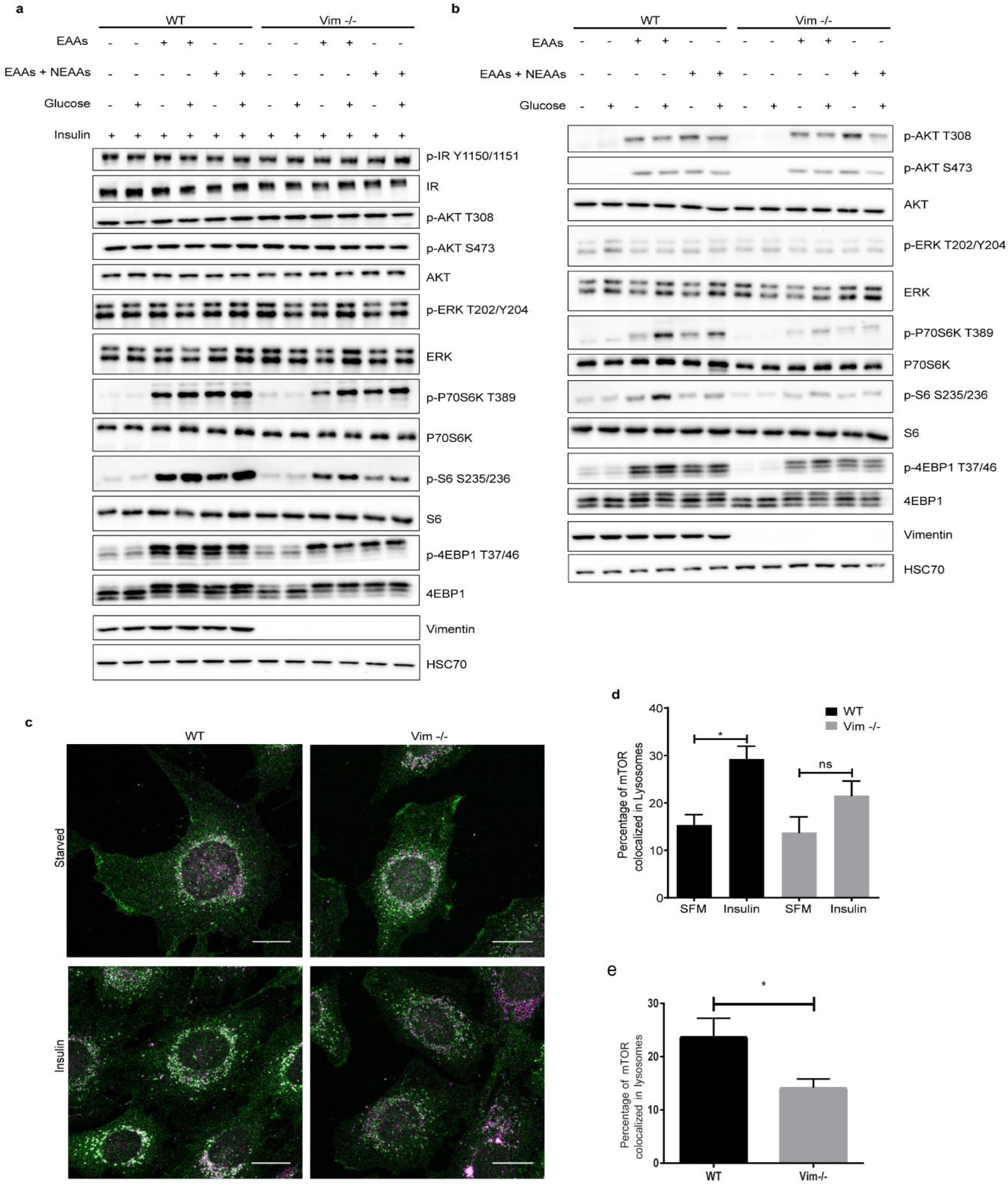
Vimentin modulates the mTORC1 pathway through nutrient and insulin signaling. **a.** Western blot analysis of WT and Vim -/- MEFs starved with RPMI media lacking amino acids, glucose and growth factors for one hour, followed by a 30 minutes stimulation with (1 mM L-glutamine plus 1x EAAs or EAAs and NEAAs) and 2.5 mM glucose plus 100 nM insulin (n=3). **b.** Same experiment as in **a.** performed without insulin. **c.** Representative images of WT and Vim -/- MEFs starved overnight and stimulated with 100 nM insulin for 15 minutes. The cells were stained for mTOR and the lysosomal marker LAMP1 (Scale bar = 20 μm). **d.** The colocalization analysis is represented as mean ± SEM of the percentage of colocalized pixels before and after insulin stimulation e. The colocalization analysis is represented as mean ± SEM of the percentage of colocalized pixels in steady state conditions and it was performed using BioImageXD (n=3). *p<0.05, ns=non-significant.

When examining the effect of nutrients alone, the overall finding is that in Vim -/- fibroblasts there is very little signaling passing down to the targets downstream of mTORC1 (Fig. 4b). While the addition of glucose to WT MEFs boosted the amino acid-mediated activation of P70S6K and the ribosomal protein S6, this amplification was basically absent in Vim -/- MEFs (Fig. 4b). Although, the S6 activation stayed overall at very low levels in the Vim -/- MEFs, some residual S6 activation could be observed, with slight variation between different nutrient treatments (Fig. 4b). However, 4EBP1 was unvaryingly inhibited in the samples from the Vim -/- MEFs (Fig. 4b). This observation may relate to the different functions of S6 and 4EBP1. 4EBP1 is an inhibitory protein, whose functions are regulated by mTORC1-mediated phosphorylation. When hypophosphorylated, 4EBP1 blocks protein synthesis by binding to the eIF4E complex. Upon phosphorylation through mTORC1, the interaction is inhibited(2,27). These results imply that in Vim -/- cells there is a consistent inhibition of protein synthesis through hypophosphorylation of 4EBP1, thereby binding the eIF4E complex.

To verify that only mTORC1 downstream signaling is affected, we performed the same experiment on cells treated with 100 nM of rapamycin, an inhibitor of mTORC1. We found that this treatment inhibits mTORC1 activation in WT MEFs and the residual mTORC1 activation in Vim -/- MEFs (Extended Data Fig. 2a, b), validating that vimentin modulates mTORC1 activation by regulating its capability to activate downstream effectors. Altogether, these results show that vimentin is required for the phosphorylation of mTORC1 downstream effectors to take place, pointing to a role of vimentin in mTORC1 activation itself.

Since the effect of vimentin was narrowed down to direct targeting of mTORC1, we wanted to examine the possibilities how this could take place. In this respect, when activated, mTORC1 translocates from the cytosol to lysosomes for activation (3,22). This is an essential step for mTORC1 activation by Rags(22). Therefore, we measured mTORC1 translocation in serum starved MEFs upon insulin stimulation using confocal imaging and colocalization analysis. While insulin stimulation of WT MEFs induced a clear colocalization of mTOR with the lysosomal marker LAMP1, no increase in the colocalization could be observed in Vim -/- MEFs (Fig. 4c, d). We also found that mTORC1 localization in lysosome is significantly reduced in Vim-/- MEFs under steady state conditions (Fig. 4e). This demonstrates mTORC1 the translocation machinery requires vimentin. This finding shows that vimentin has the ability to support mTORC1 localization to the lysosomal membrane. This modus operandi would is in agreement with the established signaling functions of vimentin, which has been identified as a scaffolding system for many signaling events and pathways (23–26).

S6 is phosphorylated at serine 234/235 through P70S6K, which is downstream of the AKT-mTORC1 signal transduction pathway(21). S6 can also be activated by P90S6K, a downstream effector of ERK signaling(28). Therefore, we used AKT and ERK kinase inhibitors to determine whether these two kinases play a role in amplifying S6 phosphorylation in a vimentin-dependent manner. We treated MEFs WT and Vim -/- with either an ERK or AKT inhibitor after 40 minutes of full starvation and performed the nutrient and insulin stimulation protocol described before. In both WT and Vim -/- MEFs, AKT inhibition completely blocked AKT and its downstream mTORC1 signaling, including phosphorylation of P70S6K, and S6 (Extended Data Fig. 2c). However, ERK inhibition had no effect on mTORC1 downstream signaling (Extended Data Fig. 2c). Thus, vimentin modulates mTORC1 downstream signaling independently of ERK signaling and the phosphorylation of S6 requires an active mTORC1.

As vimentin has been shown to regulate autophagy(15), we wanted to determine whether the results we obtained with mTORC1 signaling could be coupled to the vimentin-mediated regulation of autophagy. During autophagy, LC3I, a cytosolic protein, is conjugated with phosphatidylethanolamine to form LC3-II, which is found on isolation membranes and autophagosomes, and to some extent on autolysosomes(29). p62 interacts with autophagy substrates and it selectively transports ubiquitinated proteins to lysosomal cargo for autophagy, a process during which p62 also gets degraded. Therefore, inhibition of autophagy will increase p62 levels, while activation of autophagic flux will reduce them(30).

Importantly, we observed that nutrient starvation significantly decreased p62 levels in Vim -/- MEFs, suggesting activation of the autophagic flux, with prominently lower p62 levels as compared to the samples from starved WT MEFs (Fig. 5a). In Vim -/- MEFs, this could be partly inhibited in cells stimulated with insulin, amino acids, and glucose (Fig. 5a). To further investigate the role of vimentin in autophagy, we subjected cells to depletion of various nutrient sources (glucose, EAAs or EAAs and NEAAs). WT MEFs appeared to be well protected against autophagic flux, as reflected by the steady levels of p62. Strikingly, in Vim -/- MEFs, the levels of p62 are significantly lower than in all the corresponding WT MEF samples, indicating that without vimentin, the cells are significantly more susceptible to autophagy (Fig. 5b). Notably, the p62 depletion data can be coupled to the inhibition of the mTORC1 pathway (Fig. 5B), similarly as shown in previous figures. To further strengthen these observations, we employed the conversion of the LC3I and LC3II as autophagy markers. We found that nutrient starvation increases the conversion of LC3I to LC3II in both WT MEFs and Vim -/- (Extended Data Fig. 3). However, in Vim -/- MEFs, the rate of conversion is higher. This suggests that Vim -/- MEFs cannot withstand limited nutrient levels and are more prone to autophagy than WT MEFs. Importantly, our results show that WT MEFs can sustain the levels of p62 even in the absence of insulin and that, conversely, Vim -/- MEFs activate autophagy even in the presence of nutrients. Thus, the inhibition of autophagy in these cells is likely to take place by mechanism different from the one previously proposed (15), in which vimentin and Beclin 1 are phosphorylated by AKT(15). This notion is especially relevant as in these experiments following nutrient starvation, we saw no difference in AKT activation. Thus, as the described mode of mTORC1 regulation is independent of AKT, the mechanism that we have observed is different from the AKT and vimentin and 14-3-3-dependent mechanism previously described (15). It is well established that mTORC1 can be activated by nutrients alone(6), and that mTORC1 inhibits autophagy by phosphorylating ULK1(8), one of the proteins required for autophagy initiation via Beclin-1 phosphorylation(9). Therefore, the increased autophagy in Vim -/- cells is likely to be due to the lower mTORC1 activation, which leads to poor inhibition of the autophagic flux. This implies that vimentin would affect autophagy through mTORC1-mediated inhibition of ULK1-Beclin (Fig. 6).

**Figure. 5.**
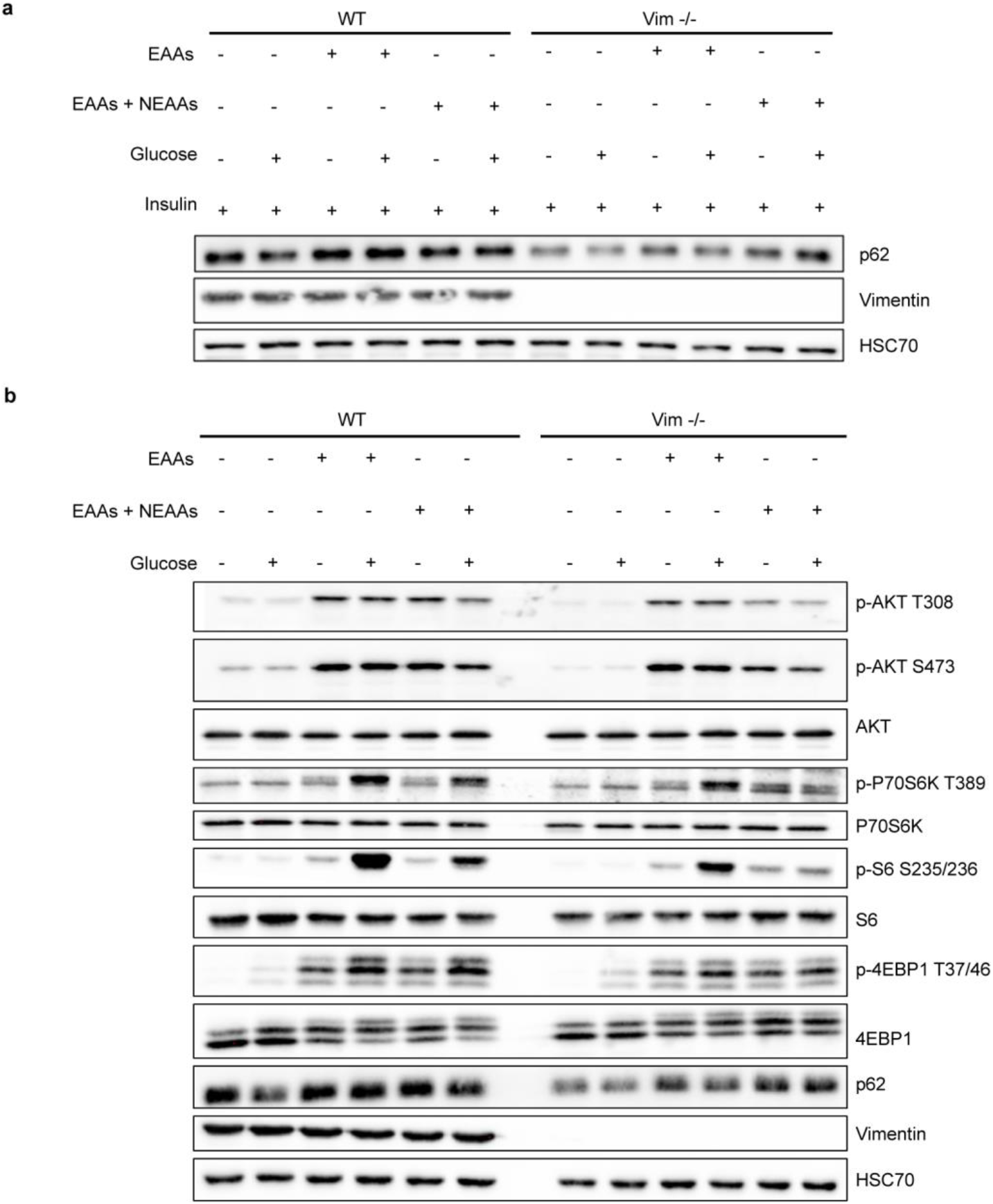
Vimentin protects MEFs from autophagy. **a.** Western blot analysis of WT and Vim -/- MEFs starved with RPMI media lacking amino acids, glucose and growth factors for one hour, followed by a 30 minutes stimulation with EAAs or EAAs and NEAAs and 2.5 mM glucose, 100 nM insulin (n=3). **b.** Western blot analysis of WT and Vim -/- MEFs grown one hour in RPMI media with different nutrients (EAAs or EAAs and NEAAs and 2.5 mM glucose; n=3).

**Figure. 6.**
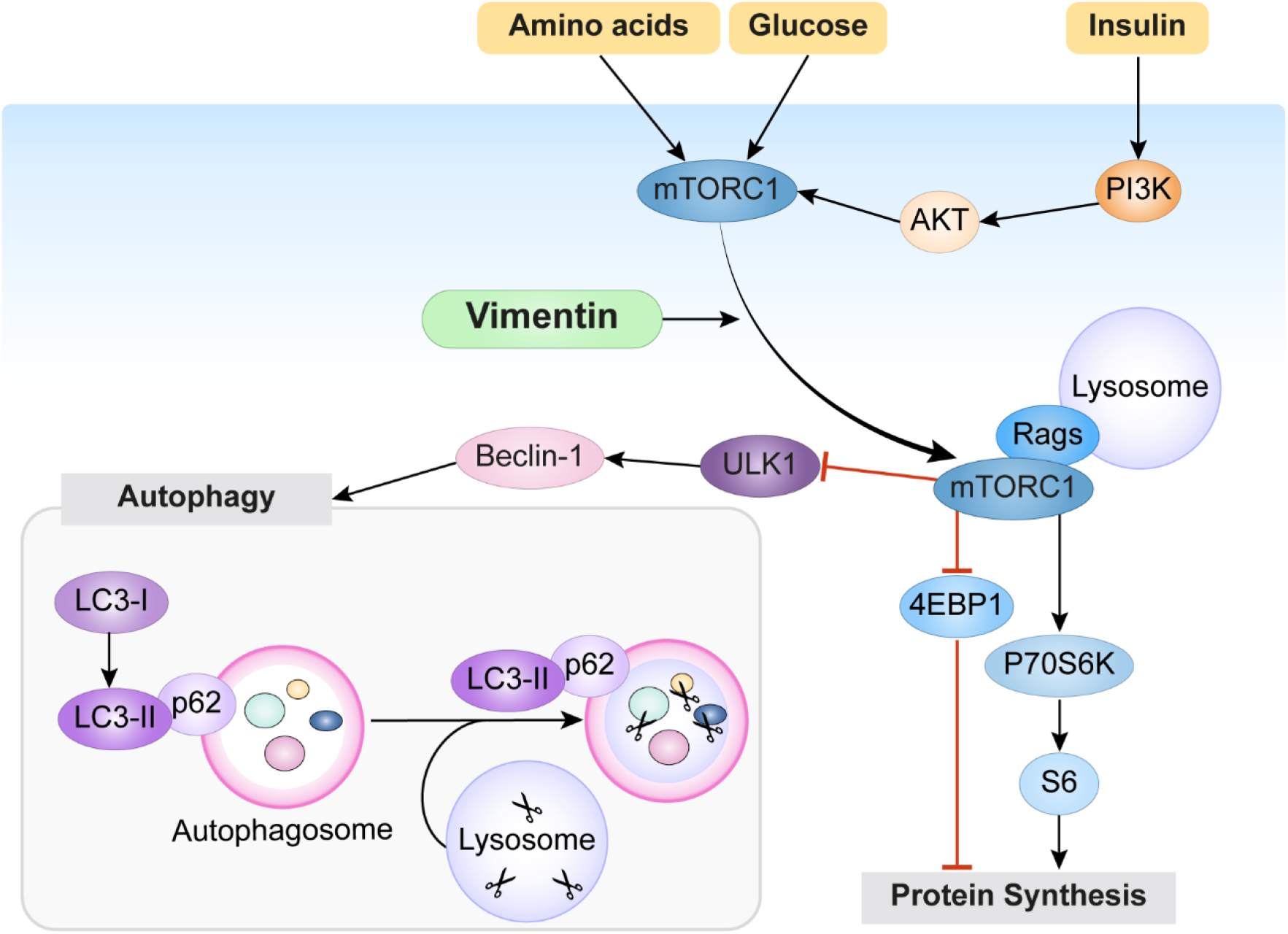
Vimentin regulates mTORC1 signaling by facilitating mTORC1 localization to the lysosome. A simplified scheme of the mTORC1 signaling system. Our results demonstrate that in the presence of nutrients, vimentin facilitates mTORC1 activation by recruiting it to the lysosomal membrane, where it can be activated by Rags. This results in the phosphorylation of 4EBP1 and P70S6K downstream targets, which lead to protein synthesis. Moreover, activation of mTORC1 results in decreased autophagy, as demonstrated by higher levels of the autophagic flux indicator p62 and reduced LC3I conversion to LC3II. As this autophagy inhibition works even under nutrient-starved condition, in the absence of Akt activation, the results imply that vimentin inhibits autophagy by mTORC1-mediated inhibition of ULK1-Beclin pathway. In this way, vimentin promotes cell size in two directions: by boosting protein synthesis through mTORC1 activation and by inhibiting of protein degradation through inhibition of autophagic flux.

Here we unravel a novel role of vimentin in cell size regulation, as the whole cell size machinery seems to be guarded and concerted by the presence of vimentin. Moreover, we provide evidence that vimentin regulates these processes by mediating mTORC1 activation. During wound healing and regeneration, the presence of nutrients and growth factors is crucial for cell growth and proliferation. Fibroblasts are rapidly proliferating cells that are crucial for the formation of granulated tissue in injury areas (31). In these dynamic but stressful conditions, vimentin levels and its organization are actively coordinated. Our results show that vimentin is the critical link between cell size signaling and the other processes critical for repair. The advantage of a vimentin-based link to cell size signaling is that vimentin is also coupled to cytoskeleton-mediated sensing of cell shape(32) and to interactions with both the surface and other cells(11). High expression of vimentin is a hallmark of many types of cancer cells. Providing an active regulation of cell size and resisting autophagy is, obviously, an additionally advantageous asset that vimentin provides for cells in dynamic transition as well as for cancer cells.

## Materials and methods

### Plasmids and antibodies

The pCMV-script-vimentin and pCMV-script empty vector plasmids were a kind gift from Professor Johanna Ivaska(33). The antibodies used for the experiments are listed in the Extended Data Table 1.

### Mice

WT and Vim -/- male mice from the strain SVJ/129 were used for all experiments. The mice were held at Central animal laboratory, Biocity unit and were fed with standard diet and free access to water. Mice were sacrificed by cervical dislocation. Genotyping was determined by polymerase chain reaction. All experiments were performed according to the guidelines set by the Ethics Committee. The body weight was measured at three (n=5) and six (n=4) months old. BMI and Lee’s index of 6 months old mice were calculated accordingly. The total fat mass was measured by EcoMRI and normalized to the body weight. To evaluate organ size and weight, WT and Vim-/- mice of 3 months of age (n=3) were sacrificed. The white adipose tissues were fixed with paraffin. The sectioning and hematoxylin-eosin staining was carried out by Lounais-Suomen pathology laboratory. The sections were imaged using the Panoramic Slide Scanner (3DHISTECH, Hungary) and Fiji was used to measure cell size.

### Cell culture and treatments

SV40-immortalised WT and Vim -/- MEFs were cultured in Dulbecco’s modified media (DMEM, Sigma #D6171) supplemented with 10 % fetal bovine serum (FBS, Biowest #S1810), 2 mM of L-glutamine (Biowest #X0550), 100 U/ml of penicillin and 100 μg/ml of streptomycin (Sigma #P0781) at 37 ºC in a 5% CO_2_ incubator and passaged when they were about 80 % confluent.

To test if the size phenotype could be rescued, Vim -/- MEFs were seeded in 6-well plates on the day prior to transfection. The cells were transfected with 7.5 μg of pCMV script-vimentin or empty vector using Xfect™ (Clontech Laboratories, USA), according to the instructions of the manufacturer. The cells were used for cell volume analysis 48 hours post-transfection. To assess how growth factors affect the cell size in MEF WT and Vim -/-, the cells were incubated in serum free media for two to three days. Then, the media was replaced with fresh serum-free media and the cells were stimulated either 100 μg/ml insulin (Sigma #I0516) or 5 % fetal calf serum (FCS, Biowest #S1710) for 24 hr. To study mTORC1 activation, 200,000 cells were seeded in 6-well plates and serum starved overnight. Next, the cells were treated either with 100 nM insulin or 5 % FCS for 15 minutes. Alternatively, the cells were fully starved for one hour with RPMI media (Life Science #R9010-01) lacking glucose, amino acids and serum. Then, the cells were treated for 30 minutes with 1 mM of L-glutamine plus 1x concentration of either minimum essential amino acids mix (EAAs)(Thermo-Scientific #11130036) or EAAs and non-essential amino acis mix (NEAAs) (1:1 mix of EAAs with NEAAs, Sigma #M7145) either in the presence or absence of 2.5 mM of glucose (Thermo scientific #A2494001) for 30 minutes. This experiment was repeated with 100 nM of insulin along with amino acids and glucose. To further investigate mTORC1 signaling, the cells were treated with 100 nM of rapamycin (Tocris #1292) for 20 minutes. This was conducted after 40 minutes of full starvation and was followed by stimulation with amino acids, insulin and glucose as described above. Alternatively, two other kinase inhibitors were used: 1 μM and 1.5 μM of AKT VIII (Calbiochem #124018) or 100 nM and 200 nM of trametinib (Selleckchem #GSK1120212). To study autophagy, 200,000 WT and Vim -/- were seeded in 6-well plates and incubated for 16 hours. On the next day, the media was replaced with RPMI media containing EAAs, with or without NEAAs and glucose for one hour. The protein levels were evaluated using immunoblotting.

### Primary MEFs isolation

The primary MEFs were isolated according to a previously described method(34). Briefly, the mice were sacrificed 13-14 days post-coitum by cervical dislocation, the uterine horns were dissected, rinsed with 70 % ethanol and kept in PBS. Under the laminar hood, the embryos were separated, and both head and red organs of the embryos were stored for genotyping. The remaining tissue was washed with PBS and minced until it became possible to pipette it. The tissues were incubated for 15 min at 37 °C with 2 ml of 0.05 % trypsin/EDTA containing 100 K units of DNaseI and the cells were dissociated by pipetting every five minutes. The trypsin was inactivated by adding about 1 volume of freshly prepared DMEM complete media and the cells were centrifuged (100 g) for 5 minutes. The supernatant was carefully removed, the cell pellet was resuspended in pre-warmed DMEM complete media and plated approximately one embryo per plate.

### Bone marrow-derived dendritic cells (BMDCs)

Primary dendritic cells were differentiated from bone marrow precursors isolated from WT and Vim -/- mice. The differentiation was performed using IMDM medium containing FCS (10 %), glutamine (20 mM), penicillin–streptomycin (100 U/ml), 2-ME (50 μM) further supplemented with granulocyte-macrophage colony-stimulating factor (50 ng/ml)-containing supernatant obtained from transfected J558 cells, as previously described(35). The semi-adherent cell fraction, corresponding to the CD86^+^ dendritic cells, was retrieved from the cell culture dishes at the differentiation days 10-12 by gentle flushing and used for the measurements. All cell culture reagents were purchased from Thermo-Scientific (Waltham, MA, USA).

### Cell area

Phase contrast images from WT and Vim -/- MEFs (approximately 200 cells in total) were taken using CellIQ (ChipMan Technologies, Finland) using a 10X objective. The cell area was manually measured using the freehand tool in Fiji(36). Cell area of WT and Vim -/- BMDCs was evaluated based on images taken for undeformed (inlet) or deformed (using two different flow rates; fr1 = 0.16 μl/s and fr2 = 0.32 μl/s) cells in a real-time deformability cytometry (RT-DC) setup(20) using 40X objective (EC-Plan-Neofluar, 40×/0.75; #420360-9900, Zeiss, Germany). The RT-DC measurements were performed according to previously established protocol(37) using a 30-μm constriction channel. The cell contours were identified using thresholded images, and the cell cross-sectional area was derived from a convex hull of the fitted contours.

### Cell volume

Immortalized and primary WT and Vim -/- MEFs were grown in 6-well plates overnight. The samples were harvested and fixed with 3 % PFA for 15 minutes and then stained with 2 μM of CellTracker™ fluorescent probes (ThermoFisher Scientific #C34552). Z-stacks were taken using a Leica TCS SP5 Matrix confocal microscope with the 20X objective. The cell volume was measured using Imaris® software 8.1 (BITPLANE, Switzerland). Three clones of MEFs were used for the analysis (approximately 2000 cells in total). Cell volume analysis was also performed in cells treated for 24 hours with 1 mM thymidine (Sigma #T1895) and in Vim -/- MEFs transfected with pCMV-script-vimentin or pCMV-script empty vector. For WT and Vim -/- BMDCs, the volume was calculated by rotation of the 2D cell contours obtained from RT-DC measurements. RT-DCD data were re-analyzed from original measurements recently published(38). The calculation was performed using ShapeOut software (ShapeOut 0.9.5; https://github.com/ZELLMECHANIK-DRESDEN/ShapeOut; Zellmechanik Dresden, Germany) according to a previously described approach^30^.

### Protein concentration

WT and Vim -/- MEFs were seeded in 6-well plates and incubated overnight. Then, the cells were trypsinised and 3.0 x 10^5^ cells were lysed for 1 hour at 4 ºC in a buffer containing 150 mM NaCl, 1 % Triton X-100, 0.2 % SDS, 50 mM Tris pH 8.0 and 0.5 % of sodium deoxycholate. The protein concentration was measured using a BCA (ThermoFisher Scientific #23227), according to the manufacturer’s instructions. The protein concentration was normalized to the number of cells.

### Immunoblotting

WT and Vim -/- MEFs (treated as described above) were washed three times with ice cold PBS and lysed with 3x Sample buffer (0.625 M Tris-HCL pH 6.8, 3 % Sodium dodecyl sulfate, 30 % glycerol, 0.015 % bromophenolblue, 3 % β- mercaptoethanol). The lysates were heated for 10 minutes at 98 ºC. The samples were separated by SDS-PAGE, transferred to nitrocellulose membranes (or methanol-activated polyvinylidene difluoride membranes for LC3) and blocked one hour with 5 % milk in TBS 0.3 % Tween20. The membranes were incubated with primary antibody overnight at 4 ºC. All primary antibodies used for western blot were diluted 1:1000 in TBS 0.3 % Tween20 3 % BSA 0.02 % sodium azide. The membranes were washed 20 minutes with TBS 0.3 % Tween20 and incubated for one hour at room temperature with secondary antibody (diluted 1:10000 in TBS 0.3 % Tween20 5 % milk). The signal was detected with enhanced chemiluminescence (Amersham #RPN2236) and a BioRad chemidoc machine.

### Immunostaining

For the mTOR translocation assay, WT and Vim -/- MEFs were seeded on top of glass coverslips and serum-starved overnight. The cells were treated with 100 nM of insulin for 15 minutes, were washed once with PBS and were fixed with 3 % PFA in PBS for 15 minutes at room temperature. Then, the cells were permeabilized with PBS 0.2 % Triton X-100 for 5 minutes and blocked with 3 % BSA in PBS 0.2 % Tween20 for one hour. The coverslips were incubated overnight in a humidified chamber with primary antibody diluted 1:200 (mTOR), 1:20 (LAMP1), and 1:2000 (vimentin) in 3 % BSA PBS 0.2 % Tween20. On the next day, the coverslips were washed three times with PBS 0.2 % Tween20, for five minutes each, and incubated with secondary antibody in 3 % BSA PBS 0.2 % Tween20 for one hour in the dark, at room temperature. The coverslips were washed as described before and dipped in Milli-Q® water once. The excess water was removed, the coverslips were mounted on the slides with mowiol® 4-88 and left to dry overnight. The imaging was performed using a Leica SP5 TCS confocal microscope with the 63X immersion oil objective numerical aperture 1.32. The colocalization analysis of LAMP1 and mTOR (n=3, 12-17 cells per trial) was carried out with the colocalization tool on BioImageXD (39).

### Protein synthesis assay

WT and Vim -/- MEFs were seeded in 96-well plates and serum-starved overnight. The assay was carried out using the Click-iT® HPG Alexa Fluor® Protein Synthesis Assay Kits (ThermoFisher Scientific #C10428). Briefly, the media was changed to methionine free media with 50 μM of a methionine analogue. The cells were treated with 100 nM of insulin for 30 minutes and were washed once with PBS. The cells were fixed with 3 % PFA in PBS and permeabilized with PBS 0,3 % Triton X-100. The Click-it reaction was prepared according to the instructions of the manufacturer and added to the cells for 30 minutes in dark. Then, the cells were washed and kept in PBS. The imaging was performed with Cell-IQ (Chip-Man Technologies, Finland) using the green channel and the 10 X objective. The image analysis (n=3, 100 cells in total) was carried out with the Cell-IQ Analyzer software by measuring the signal intensity.

### Statistical analysis

All the statistical analysis was carried out with GraphPad Prism 7 (GraphPad Software Inc., USA). Three independent trials were carried out for each experiment unless it is stated otherwise. The statistical significance between two groups were measured by unpaired student t-test with Welch correction. The Mann-Whitney test was used for samples not following a normal distribution. Multiple comparisons were performed using Welch ANOVA test with Holm-Sidak correction for multiple testing. For BMDC measurements five independent experiment replicates were performed, and the statistical analysis was performed using linear mixed effect model in ShapeOut according to previously described procedures(40).

## Acknowledgments

This study was supported by Academy of Finland, Sigrid Jusélius Foundation, Magnus Ehrnrooth Foundation, the Endowment of the Åbo Akademi University, K. Albin Johanssons stiftelse, Maud Kuistila Memorial Foundation, Liv och Hälsa Foundation, Otto A Malm Foundation, Finnish Cultural Foundation, and the Foundation of King Gustaf V and Queen Victoria (“Konung Gustaf V:s och Drottning Victorias Frimurarestiftelse”). We also like to thank the Cell Imaging Core at Turku Bioscience and Biocenter Finland for the research facilities. FL thanks the DFG for financial support within the CRC 1027.

## Author contributions

Experimental design was prepared by **PM,** and **JEE**; mice experiments were performed by **FC, PM and LSCR**. MEF experiments and related data analysis were performed by **PM** and **LSCR** imaging and image analysis was performed by **PM**, **LSCR** and **MM**; BMDC experiments and related data analysis were performed by **MU** under supervision of **FL**; the manuscript was written by **PM, LSCR,** and **JEE**. All authors revised and edited the manuscript. **JEE** acquired funding.

## Supplementary data

**Extended Data Video 1:** Video on the effect of vimentin on cell size. Live cell imaging shows that Vim -/- MEFs are consistently smaller than WT MEFs grown to confluency.

**Extended Data Figure. 1.**
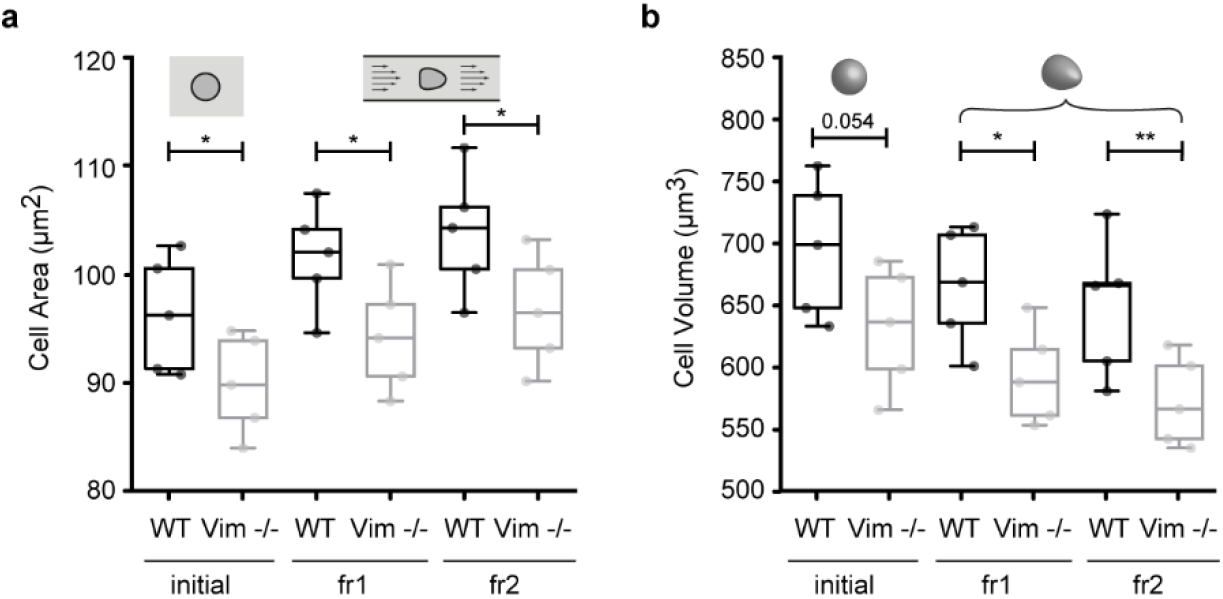
The size phenotype is not restricted to MEFs. **a.** Cell area of WT (black) and Vim -/- (gray) BMDCs obtained from RT-DC measurements of undeformed, spherical cells (initial) and cells deformed in a narrow constriction of a microfluidic channel using two different flow rates (fr1 = 0.16 μl/s and fr2 = 0.32 μl/s). **b**. Cell volume of WT (black) and Vim -/- (gray) BMDCs corresponding to the samples in **a**. In **a** and **b** each data point represents a mean of an independent RT-DC measurement (n = 5). *p<0.05, **p<0.01. RT-DC cell size data has been obtained by reanalyzing measurements recently published.

**Extended Data Figure. 2.**
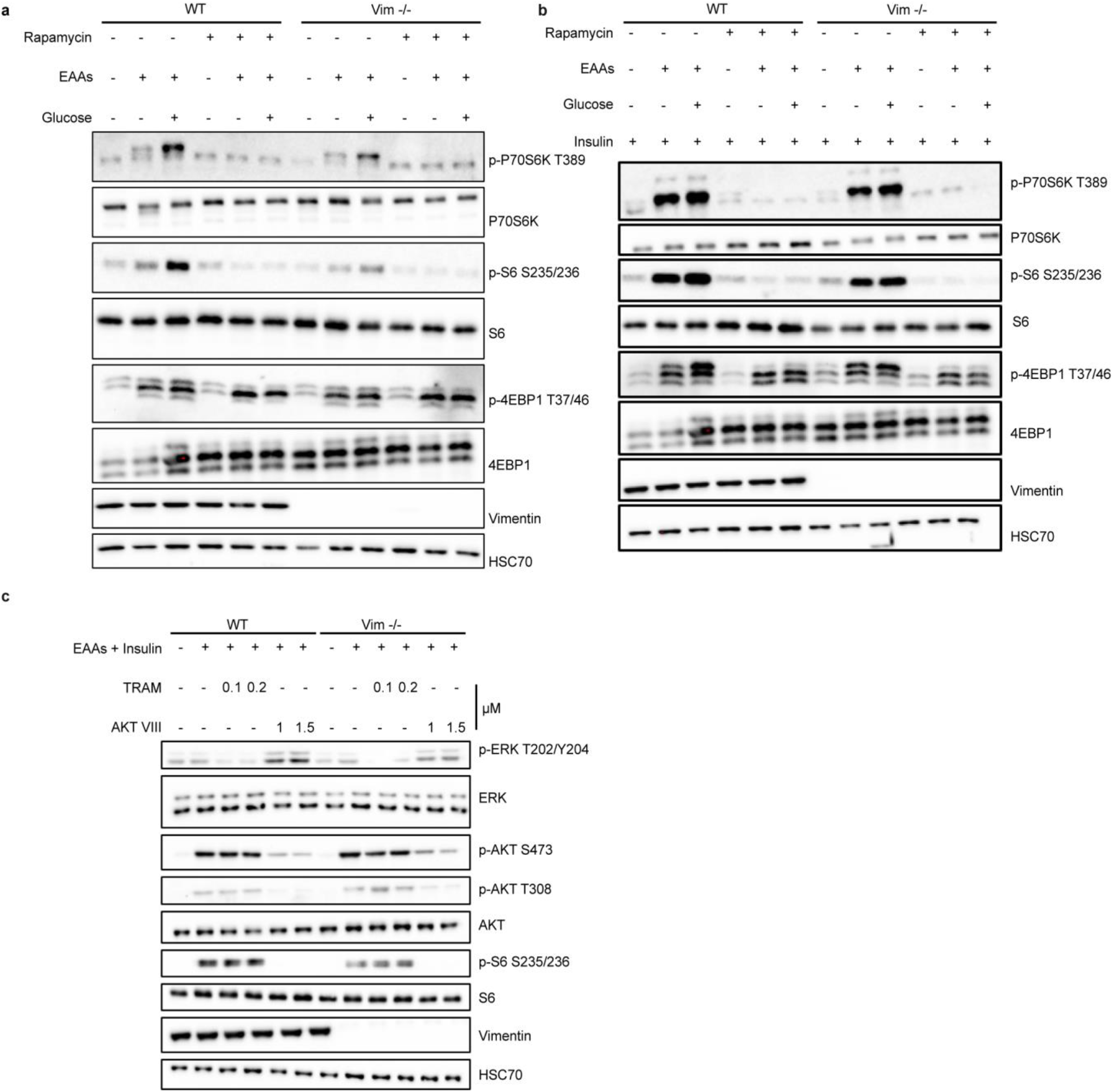
Phosphorylation of mTORC1 downstream targets requires an active mTORC1. **a.** Western blot analysis of WT and Vim -/- MEFs starved with RPMI media lacking amino acids, glucose and growth factors for 1 hour, followed by stimulation with EAAs or EAAs and NEAAs, with or without 2.5 mM glucose in all combinations. Cells were treated with 100 nM of rapamycin after 40 minutes of starvation (n=3). **b.** Same experiment but including a 100 nM insulin treatment (n=3). **c.** Western blot analysis of WT and Vim -/- MEFs treated as in Fig. 3b in the presence of an ERK (TRAM) or an AKT (AKT VIII) inhibitor (n=3).

**Extended Data Figure. 3.**
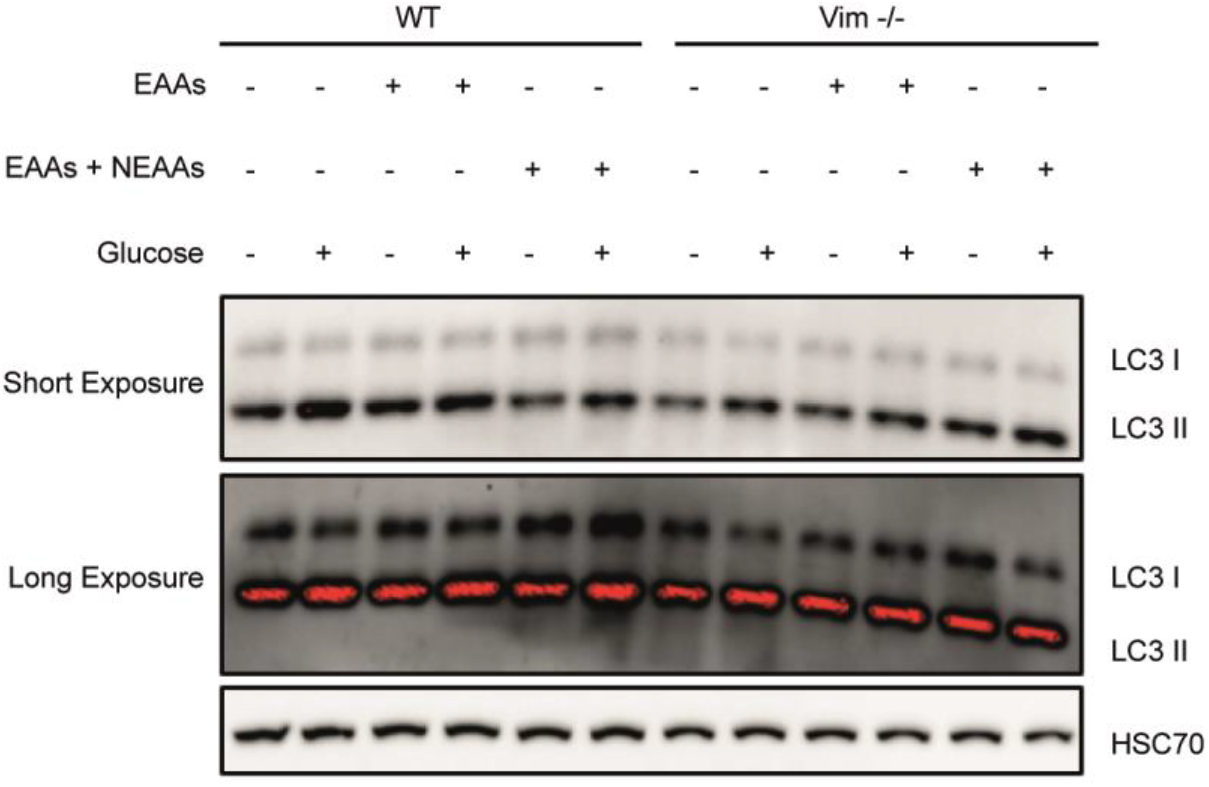
Same experiment as in Fig. 4b. to assess the level of LC3I and LC3II as an indicator of autophagy (n=3).

**Extended Data Table 1.** List of antibodies.

| Application | Primary antibody | Product details | Secondary antibody | Product details |
| --- | --- | --- | --- | --- |
| **Immunostaining** | Vimentin | Biolegend (#919101) | Chicken 555 | Thermo (#A21437) |
|  | mTOR | CST (#2983) | Rabbit 488 | Thermo (#A11034) |
|  | LAMP1 | SCBT (#19992) | Rat 633 | Thermo (#A21247) |
| **Immunoblotting** | HSC70 | Enzo (#ADI-SPA-815) | Rat HRP | GE (#NA935) |
|  | LC3 | Novus (#100-2220) | Rabbit HRP | Promega (#W401B) |
|  | P70S6K | CST (#2708) |  |  |
|  | S6 | CST (#2217) |  |  |
|  | 4EBP1 | CST (#9644) |  |  |
|  | P62 | CST (#5114) |  |  |
|  | AKT | CST (#9272) |  |  |
|  | ERK | CST (#4695) |  |  |
|  | Insulin Receptor | CST (#3025) |  |  |
|  | P70S6K Thr389 | CST (#97596) |  |  |
|  | S6 Ser235/236 | CST (#4858) |  |  |
|  | 4EBP1 Thr37/46 | CST (#2855) |  |  |
|  | AKT Thr308 | CST (#13038) |  |  |
|  | ERK Thr202/204 | CST (#4370) |  |  |
|  | AKT Ser473 | CST (#4060) |  |  |
|  | Insulin Receptor Tyr1150/1151 | CST (#3024) |  |  |
|  | Vimentin | CST (#5741) |  |  |

## Notes

https://youtu.be/O__mbLL0i10

### Competing Interest Statement

The authors have declared no competing interest.

